## supplemental figures for "Nucleolar Reorganization After Cellular Stress is Orchestrated by SMN Shuttling between Nuclear Compartments"

### SUPPLEMENTARY INFORMATION

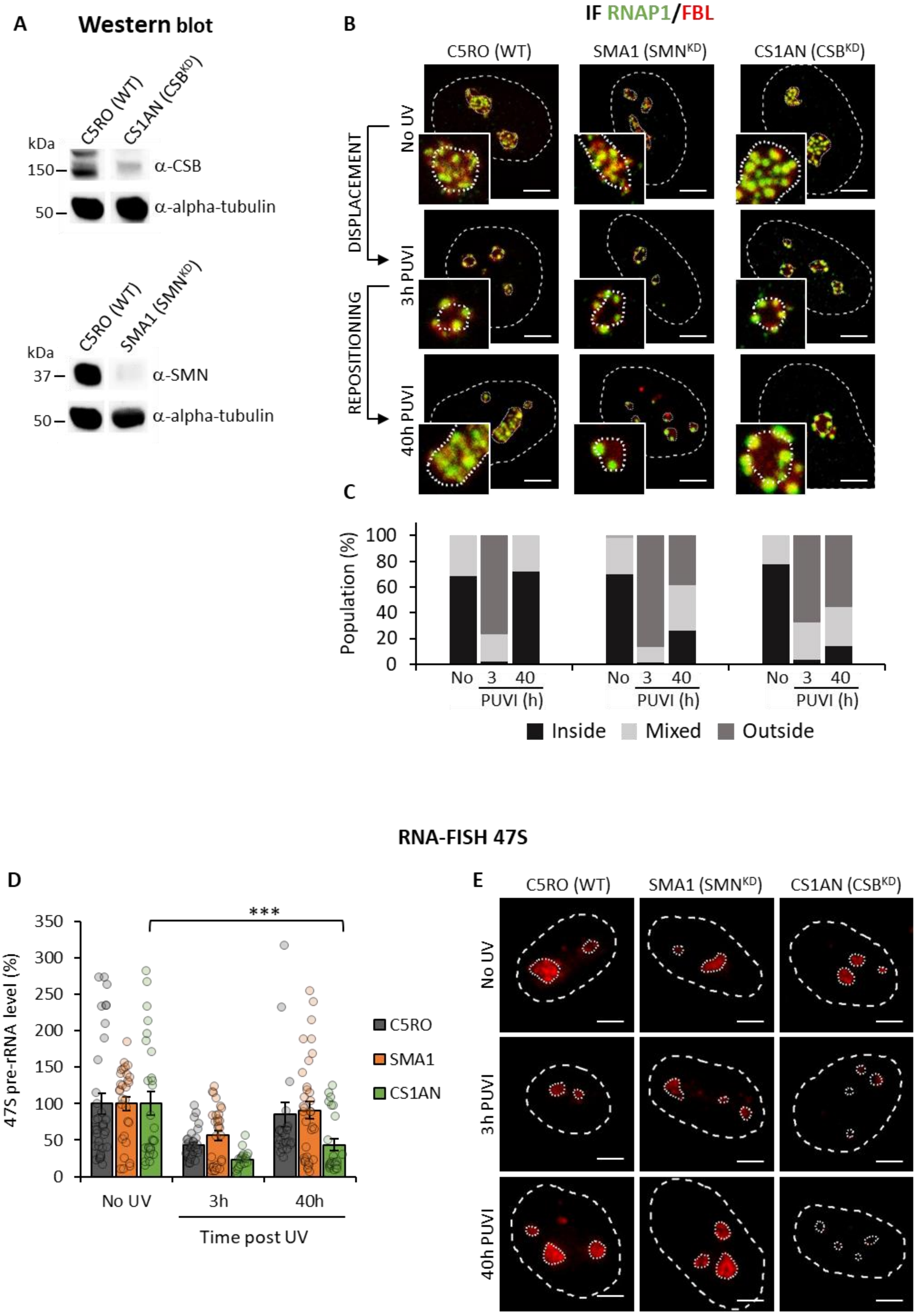
A
Western blot
C5RO (WT)
CS1AN (CSB^KD)
kDa
150
50
α-CSB
α-alpha-tubulin
C5RO (WT)
SMA1 (SMN^KD)
kDa
37
50
α-SMN
α-alpha-tubulin
B
IF RNAP1/FBL
C5RO (WT)
SMA1 (SMN^KD)
CS1AN (CSB^KD)
DISPLACEMENT
REPOSITIONING
No UV
3h PUVI
40h PUVI
C
Population (%)
No 3 40
PUVI (h)
Inside
Mixed
Outside
RNA-FISH 47S
D
47S pre-rRNA level (%)
***
No UV
3h
40h
Time post UV
C5RO
SMA1
CS1AN
E
C5RO (WT)
SMA1 (SMN^KD)
CS1AN (CSB^KD)
No UV
3h PUVI
40h PUVI

5

**Figure S1: RNAP1 and FBL localization during DNA repair in SMN deficient transformed fibroblasts.**

***(A)*** *Western Blot of SMN and CSB on whole cell extracts of primary fibroblasts.* ***(B)*** *Representative confocal microscopy images of immunofluorescence (IF) assay against RNAP1 (green) and FBL (red) in primary fibroblasts, at different times post UV-irradiation (PUVI). Nuclei and nucleoli are indicated by dashed lines and dotted lines respectively Scale bar: 5µm.* ***(C)*** *Quantification of cell number for RNAP1 localization (inside the nucleolus, outside the nucleolus or mixed localization) at different times PUVI.* ***(D)*** *Quantification of RNA-FISH assay showing the 47S pre-rRNA level after UV-C exposure in transformed fibroblasts. Error bars represent the SEM obtained from at least 27 cells. P-value of student's test compared to No UV condition: ***<0.001.* ***(E)*** *Representative images of RNA-FISH 47S in primary fibroblasts from Figure S1D. Scale bar 5µm*

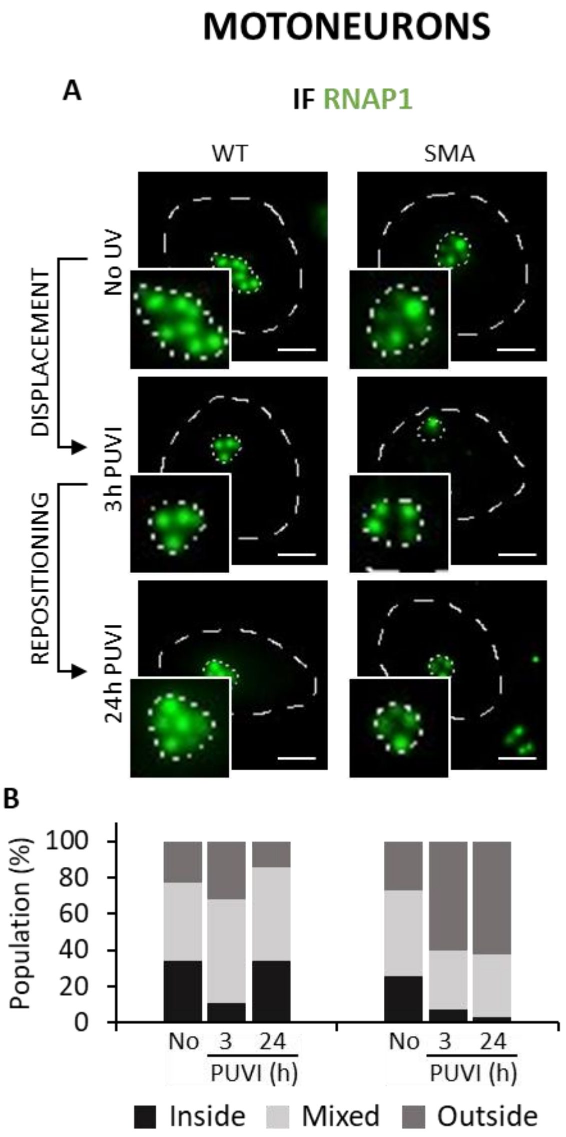


**Figure S2: RNAP1 localization during DNA repair in iPSC-derived motoneurons.**
***(A)*** *Representative microscopy images of immunofluorescence (IF) assay against RNAP1 (green) in iPSC-derived motoneurons from control (WT) or SMA patients, at different times PUVI. Nuclei and nucleoli are indicated by dashed lines and dotted lines respectively Scale bar: 5µm.* ***(B)*** *Quantification of the number of cells for RNAP1 localization (inside the nucleolus, outside the nucleolus or mixed localization) at different times PUVI.*

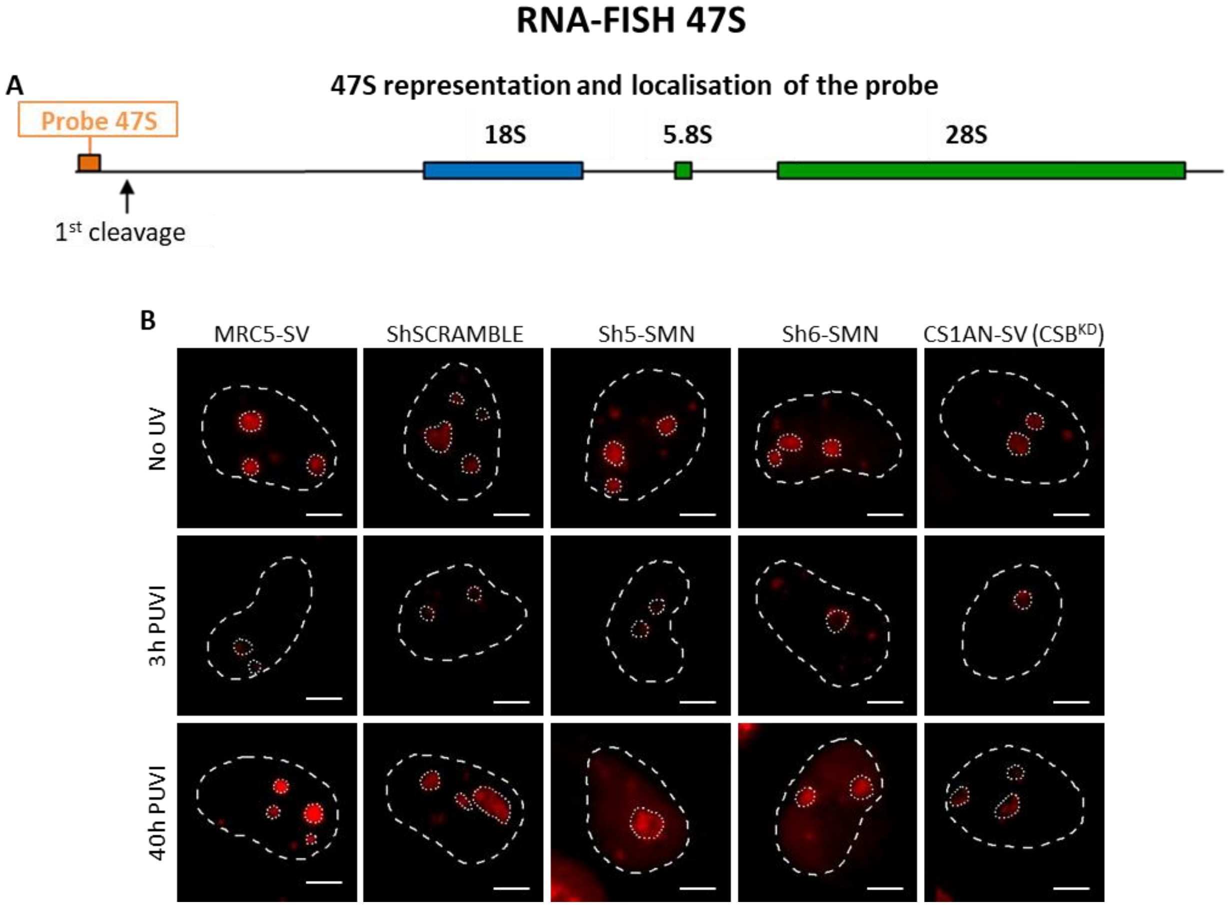


**Figure S3: RNA-FISH 47S in transformed fibroblasts.**

***(A)*** *Schematic representation of rRNA unit and localization of the 47S pre-RNA probe.* ***(B)*** *Representative images of RNA-FISH 47S in transformed fibroblasts from Figure 1D. Scale bar 5µm.*

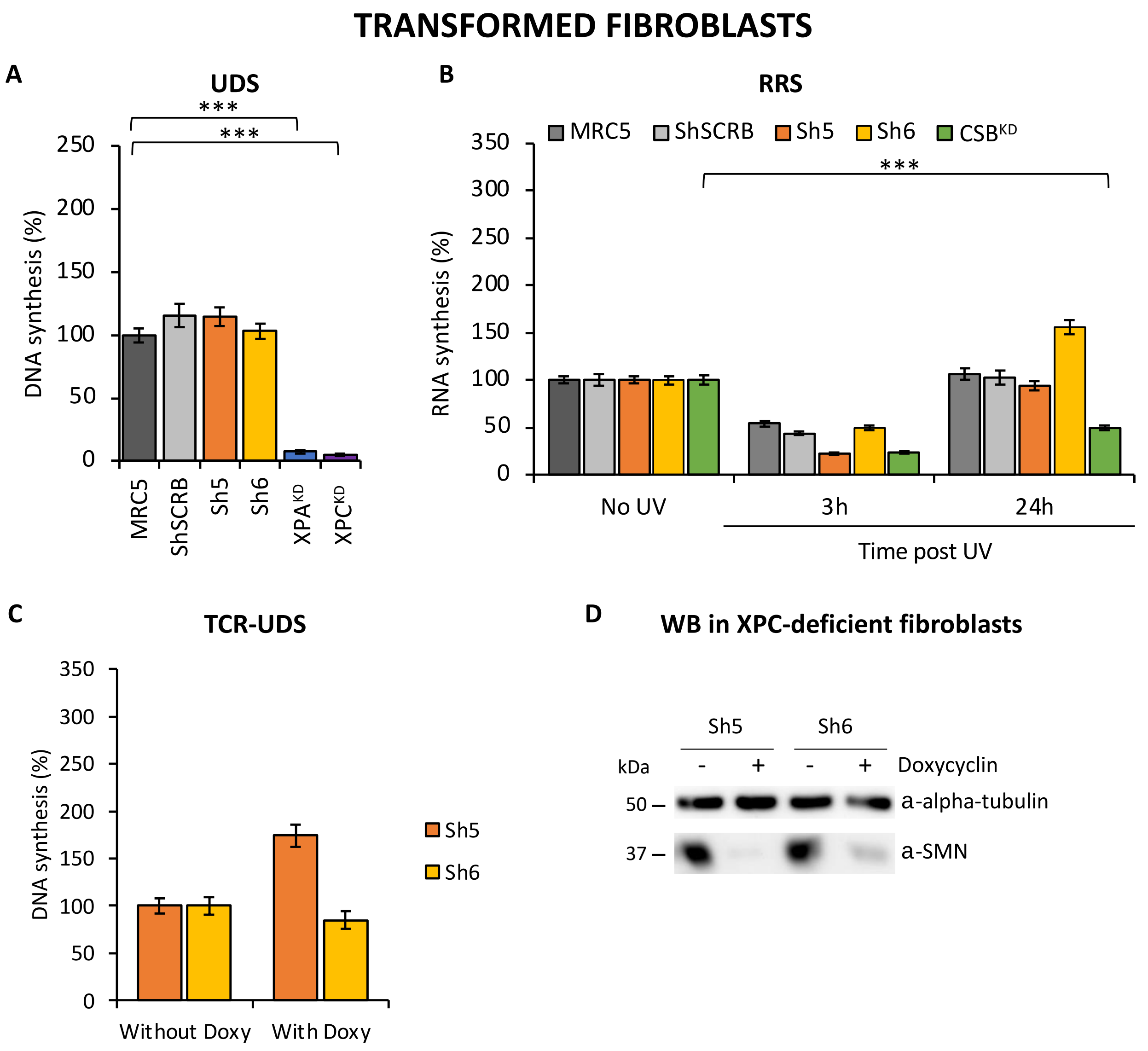


**Figure S4: SMN cells are proficient in NER**

***(A)*** *Quantification of Unscheduled DNA Synthesis assay (UDS) determined by EdU incorporation after local damage (LD) induction with UV-C (100J/m²) in transformed fibroblasts. Error bars represent the SEM obtained from at least 30 LDs.* ***(B)*** *Quantification of RNA Recovery Synthesis (RRS) assay determined by EU incorporation after UV-C (10J/m²) exposure in transformed fibroblasts. Error bars represent the SEM obtained from at least 50 cells.* ***(C)*** *Quantification of TCR-UDS assay determined by EdU incorporation after LD induction with UV-C (100J/m²) in GG-NER deficient cells (XPC-/- cells) expressing or not Sh5-SMN or Sh6-SMN. Error bars represent the SEM obtained from at least 15 LDs.* ***(D)*** *Western Blot of SMN on whole cell extracts of XPC-deficient cells with sh5-SMN or sh6-SMN. Doxycycline treatment induces the expression of the shRNA against SMN.*

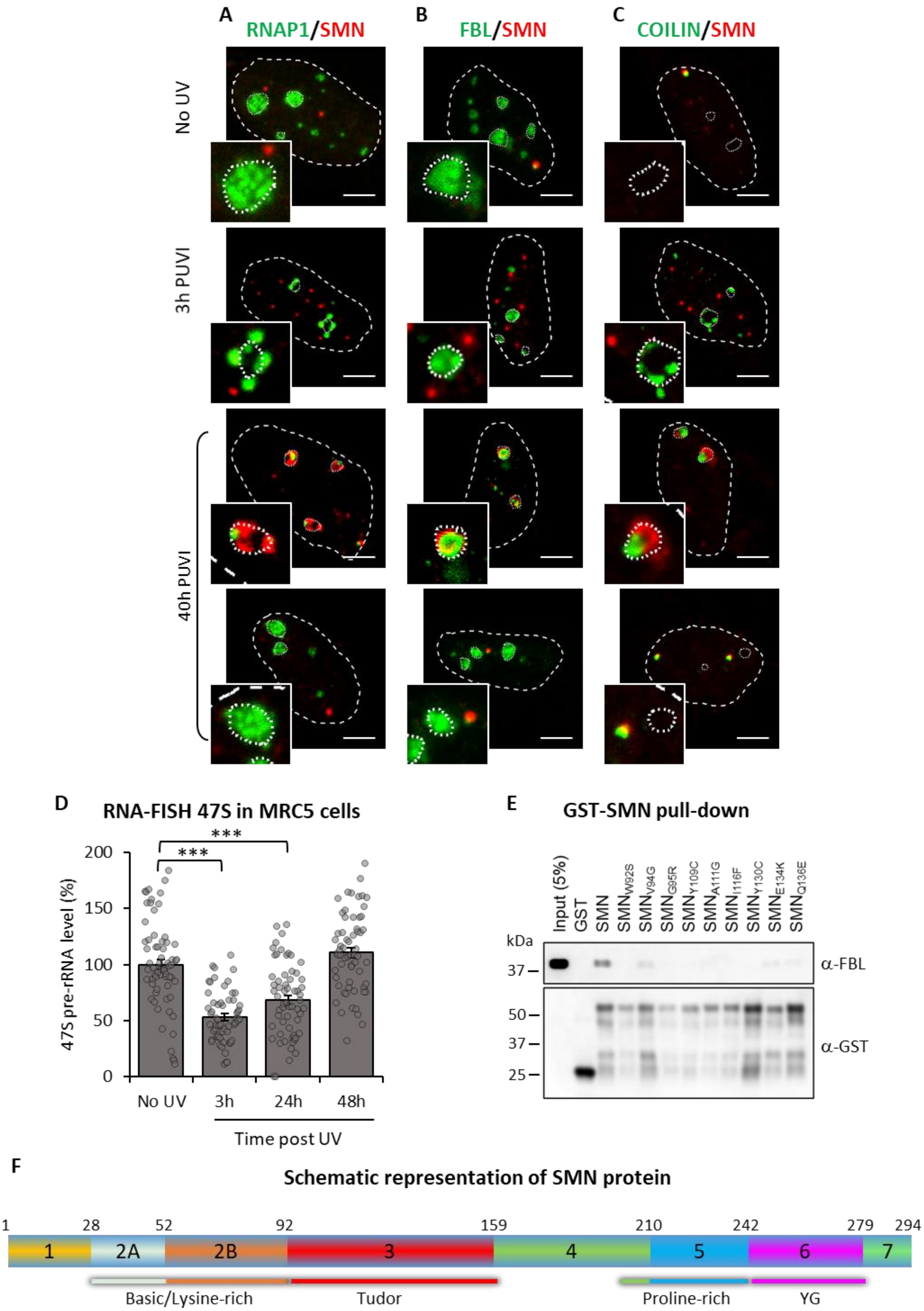

A
RNAP1/SMN
B
FBL/SMN
C
COILIN/SMN
No UV
3h PUVI
40h PUVI
D
RNA-FISH 47S in MRC5 cells
47S pre-rRNA level (%)
***
***
200
150
100
50
0
No UV
3h
24h
48h
Time post UV
E
GST-SMN pull-down
Input (5%)
GST
SMN
SMNW92S
SMNV94G
SMNG95R
SMNY109C
SMNA111G
SMNI116F
SMNY130C
SMNE134K
SMNQ136E
kDa
37
α-FBL
50
37
25
α-GST
F
Schematic representation of SMN protein
1
28
52
92
159
210
242
279
294
1
2A
2B
3
4
5
6
7
Basic/Lysine-rich
Tudor
Proline-rich
YG

**Figure S5: SMN and its partners during DNA repair**

*Representative microscopy images of immunofluorescence (IF) assay in MRC5 cells showing, after 16J/*$m^2$ *UV-C irradiation, the localization of SMN (red) and* ***(A)*** *RNAP1,* ***(B)*** *FBL or* ***(C)*** *Coilin (green) at different times PUVI. Nuclei and nucleoli are indicated by dashed lines and dotted lines respectively. Scale bar represents 5 µm.* ***(D)*** *Quantification of RNA-FISH assay showing the 47S pre-rRNA level in MRC5-SV cells after 3h, 24h and 48h of UV-C exposure (16J/*$m^2$*). Error bars represent the SEM obtained from at least 50 cells.* ***(E)*** *GST pull-down assay using purified recombinant SMN protein with different mutations in the Tudor domain found in SMA patients and cellular extracts.* ***(F)*** *Schematic representation of SMN protein with the different exons as well as domains they encode. The number of amino acids encoded by each exon is indicated.*

**A** **KBr03 treatment**

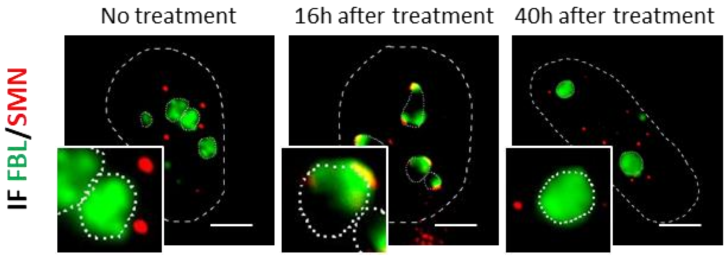


**B** **Cordycepin experiment**

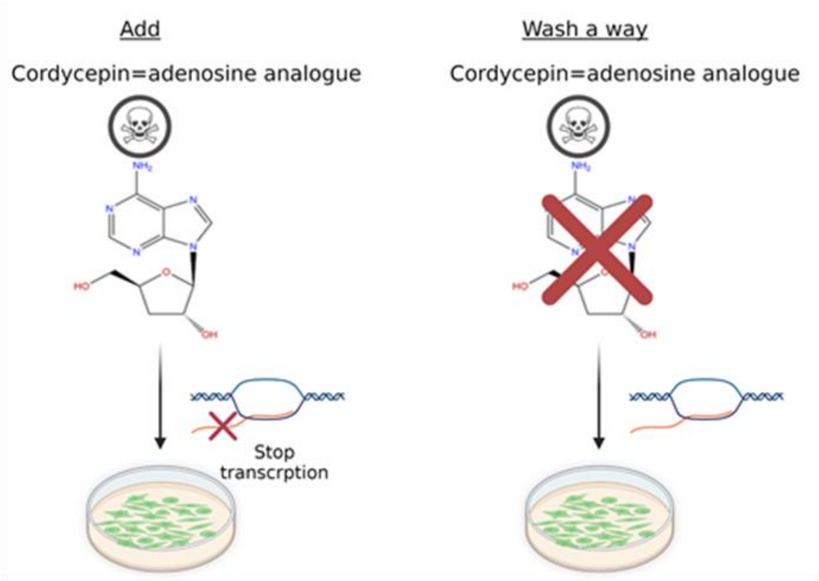


**C** **Immunofluorescence after Cordycepin treatment**

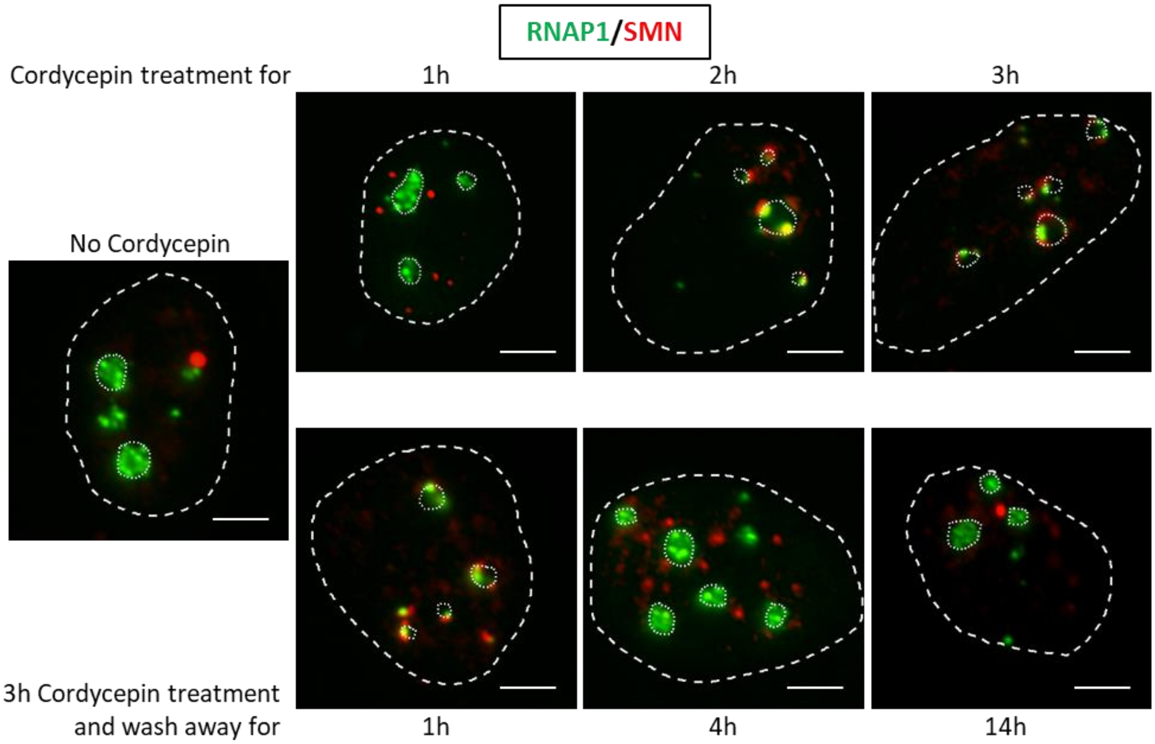

**Figure S6: SMN shuttles to the nucleolus after oxidative damage and transcription blockage**

***(A)*** *Representative microscopy images of immunofluorescence (IF) assay against RNAP1 (green) and SMN (red) after 1h of treatment with 80mM KBrO3. Nuclei and nucleoli are indicated by dashed lines and dotted lines respectively. Scale bar represents 5µm.* ***(B)*** *Schematic representation of experiment using Cordycepin.* ***(C)*** *Representative microscopy images of IF against RNAP1 (green) and SMN (red) after treatment with 50µg/ml of Cordycepin. Nuclei and nucleoli are indicated by dashed lines and dotted lines respectively. Scale bar represents 5µm.*

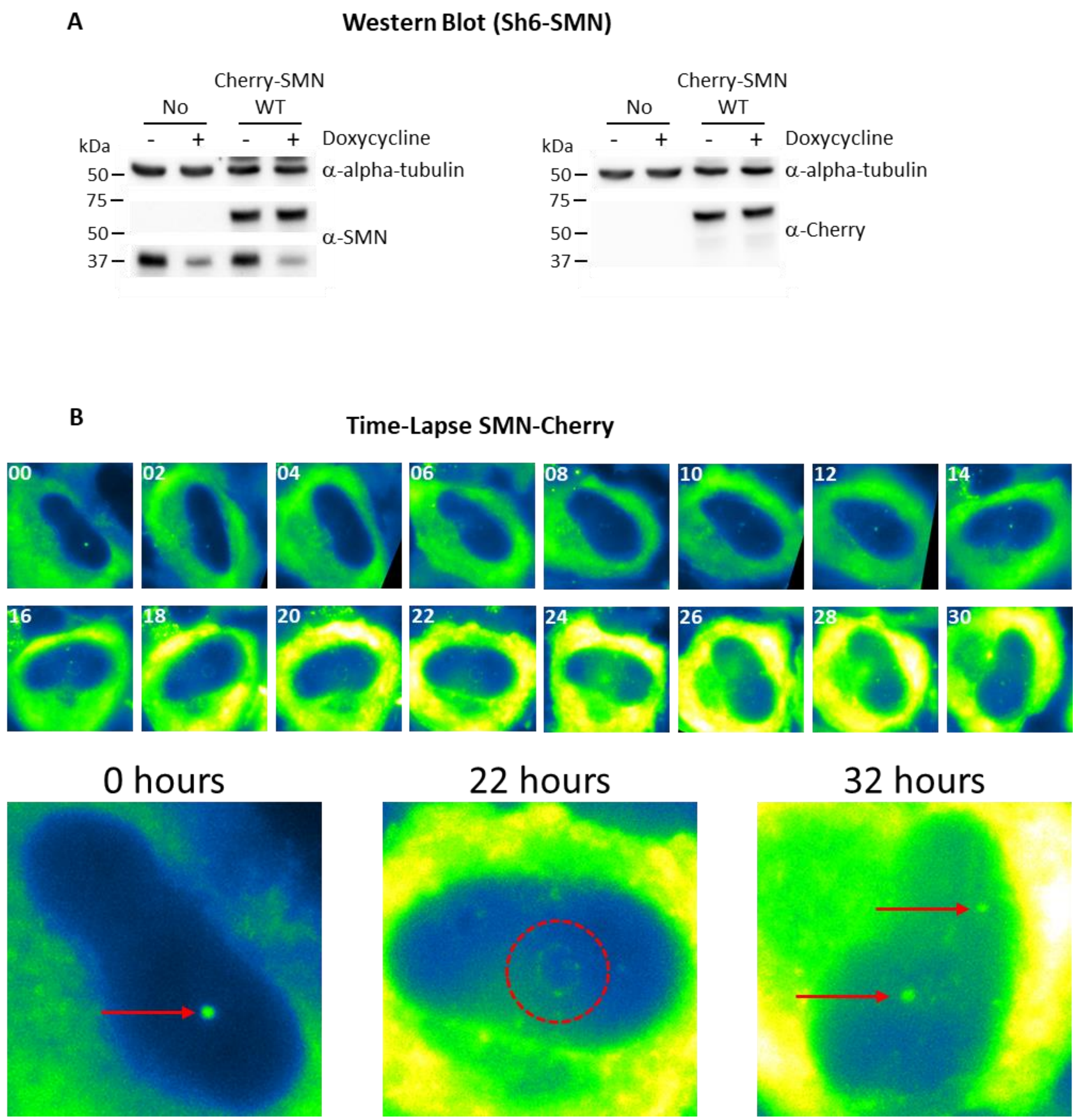


**Figure S7: SMN Shuttle to the nucleolus after UV irradiation**

***(A)*** *Western Blot of SMN on whole cell extracts of MRC-SV + Sh6-SMN transfected or not with Cherry-SMN. Doxycycline treatment induce the expression of the ShRNA against SMN.* ***(B)*** *Representative image of Cherry-SMN expressing cell, cultured in presence of Doxycycline, irradiated with 16J/m$^2$ UV-C and imaged every hour.*

### Interaction with FIBRILLARIN after UV irradiation

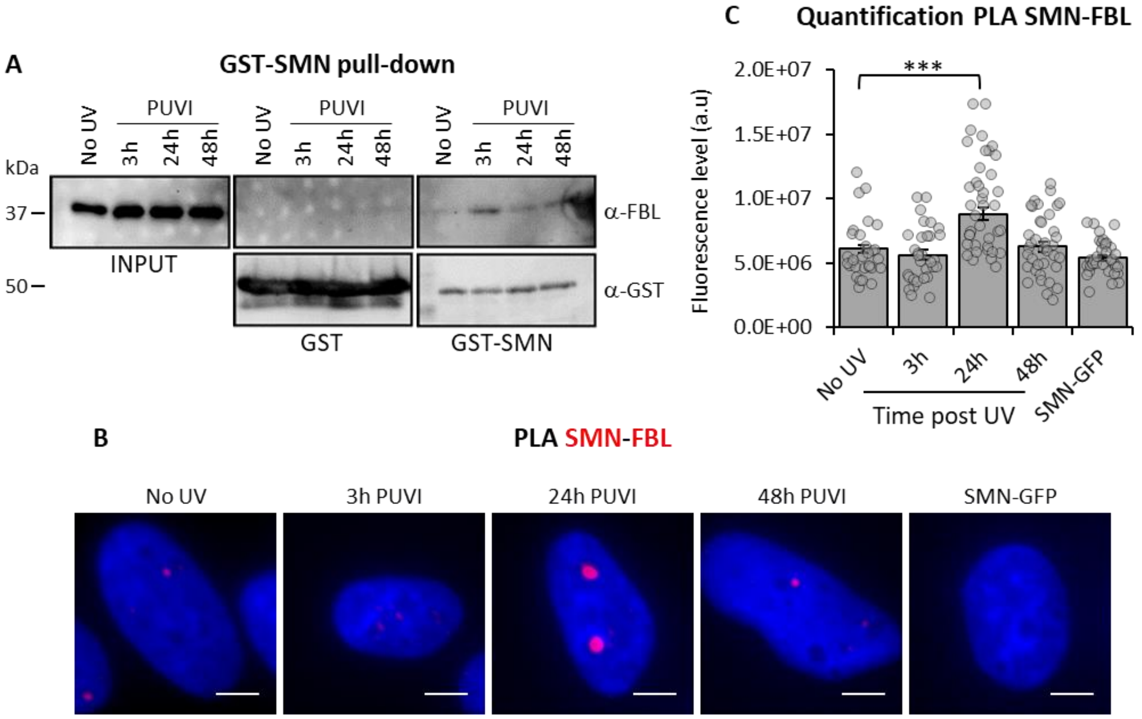


### Interaction with COILIN after UV irradiation

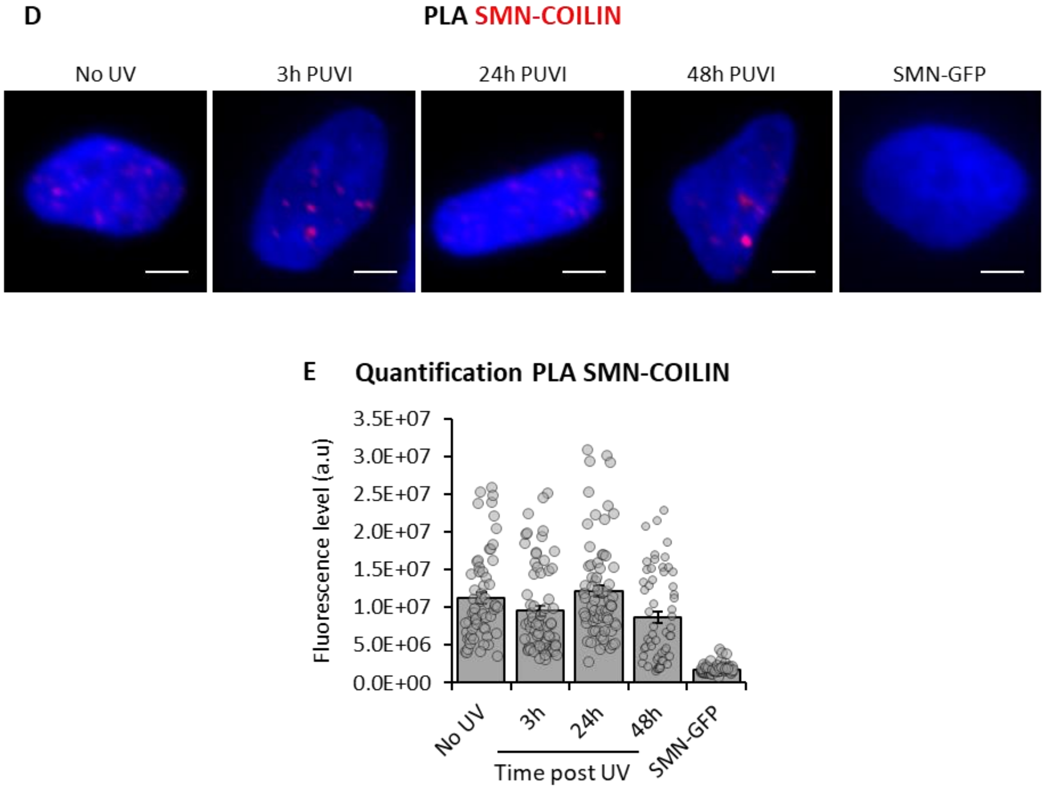

**Figure S8: SMN interacts with FBL after UV irradiation.**

***(A)*** *GST pull-down assay using purified recombinant SMN protein and cellular extracts after UV-C irradiation.* ***(B)*** *Representative microscopy images of proximity ligation assay (PLA) showing the interaction between FBL and SMN in WT cells after UV-C irradiation. Scale bar: 5µm.* ***(C)*** *Quantification of fluorescent signal in the nucleus against the couple SMN-FBL from PLA experiment in WT cells after UV-C irradiation. P-value of Student's test compared to No UV condition. ***<0.001.* ***(D)*** *Representative microscopy images of proximity ligation assay (PLA) showing the interaction between COILIN and SMN in WT cells after UV-C irradiation. Scale bar: 5µm.* ***(E)*** *Quantification of fluorescent signal in the nucleus against the couple SMN-COILIN from PLA experiment in WT cells after UV-C irradiation.*

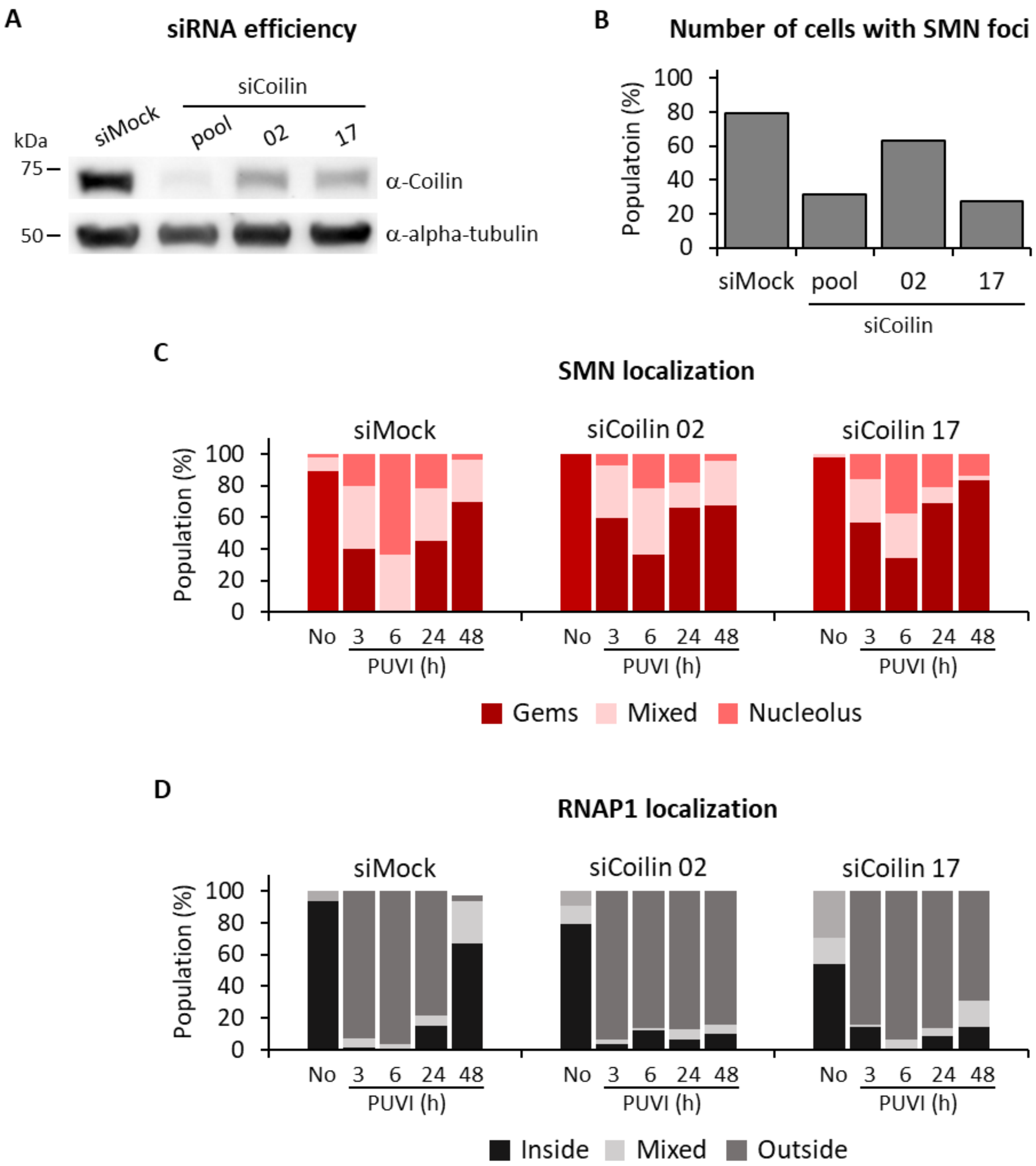


**Figure S9: SMN shuttling is Coilin-dependent**

*(A)* *Western Blot on whole cell extracts of MRC5 cells treated with siCoilin pool or individual.* ***(B)*** *Quantification of cells number with SMN foci in cells transfected with siMock or siCoilin pool or individual in No UV condition* ***(C and D)*** *Quantification of cells number for localization of* ***(C)*** *SMN (in Cajal Bodies [CBs] or Gems, at the periphery of the nucleolus or mixed localization) and* ***(D)*** *RNAP1 (inside the nucleolus, outside the nucleolus or mixed localization) at different times post UV-irradiation in cells transfected with siMock or individual siCoilin.*

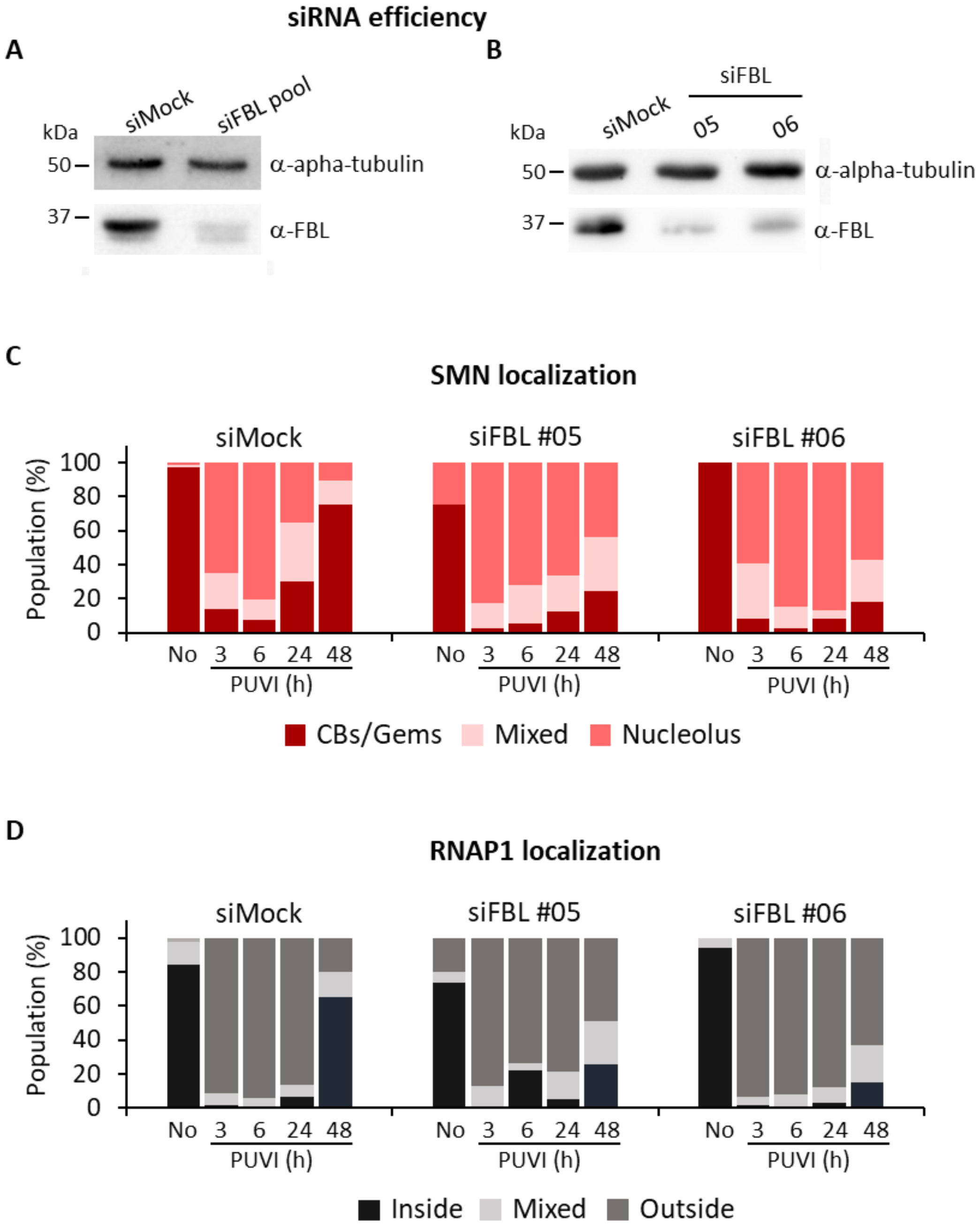


**Figure S10: FBL is required for nucleolar rearrangement during DNA repair**

***(A and B)*** *Western Blot on whole cell extracts of MRC5 cells treated with siFBL pool* ***(A)*** *or individual.* ***(B).*** ***(C and D)*** *Quantification of cells number for localization of* ***(C)*** *SMN (in Cajal Bodies [CBs] or Gems, at the periphery of the nucleolus or mixed localization) and* ***(D)*** *RNAP1 (inside the nucleolus, outside the nucleolus or mixed localization) at different times PUVI in cells transfected with siMock or siFBL individual.*

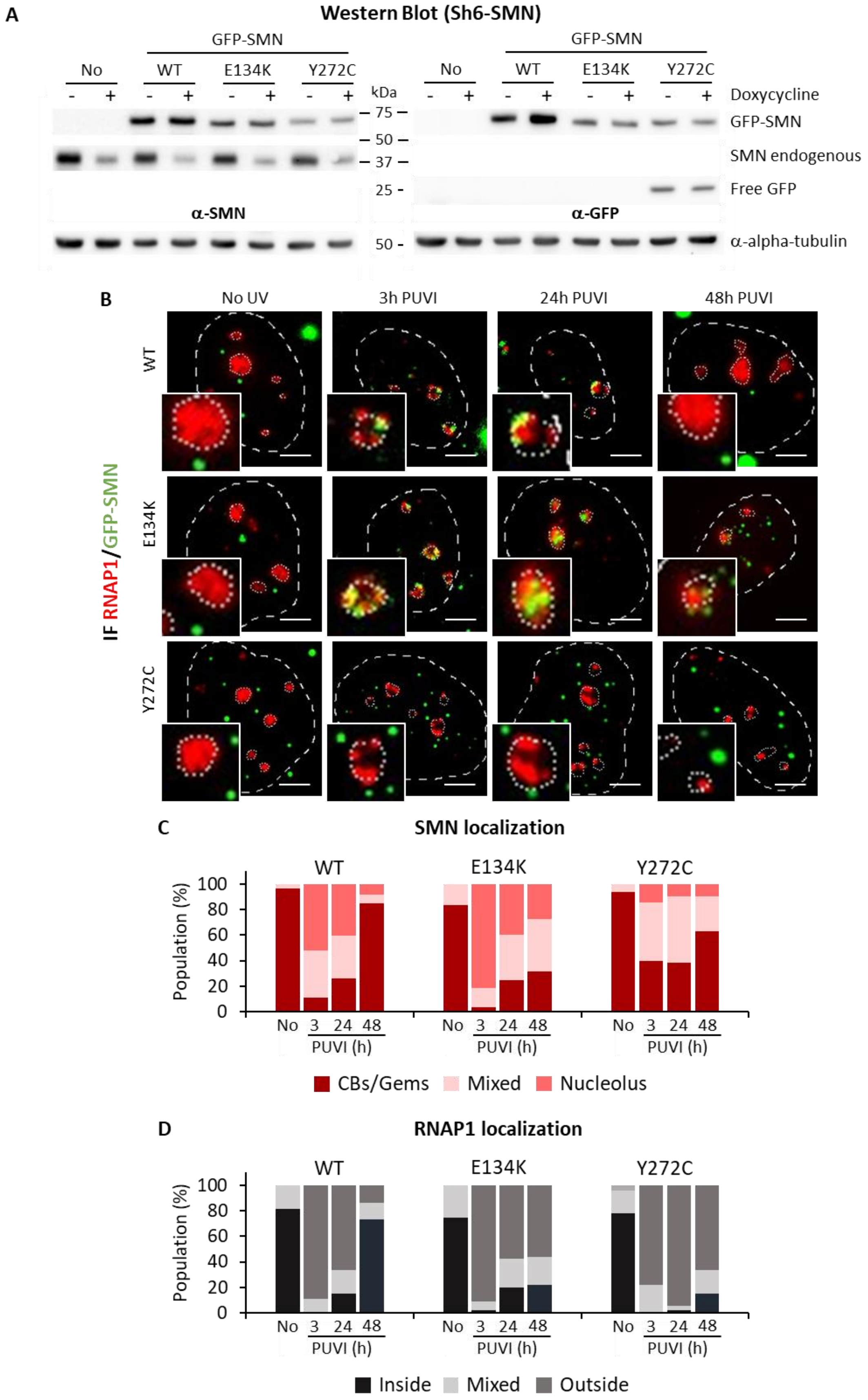

A
Western Blot (Sh6-SMN)
GFP-SMN
No
WT
E134K
Y272C
- + - + - + - +
kDa
Doxycycline
GFP-SMN
SMN endogenous
Free GFP
α-SMN
α-GFP
α-alpha-tubulin
75
50
37
25
50
B
No UV
3h PUVI
24h PUVI
48h PUVI
WT
E134K
Y272C
IF RNAP1/GFP-SMN
C
SMN localization
Population (%)
No 3 24 48
PUVI (h)
CBs/Gems
Mixed
Nucleolus
D
RNAP1 localization
Inside
Mixed
Outside


102

**Figure S11: SMN mutants are deficient in nucleolar reorganization during DNA repair.** ***(A)*** *Western Blot on whole cell extracts of MRC5 cells transfected or not with GFP-SMN, GFP-SMN*$^{E134K}$ *or GFP-SMN*$^{Y272C}$*. GFP-SMN is revealed with either SMN (left panel) or GFP (right panel) antibody. Doxycycline treatment induces the expression of the shRNA against SMN.* ***(B)*** *Representative microscopy images of IF assay showing the localization of RNAP1 (red) and GFP-SMN (green) WT (top panel) E134K (middle panel) or Y272C (bottom panel) at different times PUVI in cells depleted of the endogenous SMN. Nuclei and nucleoli are indicated by dashed lines and dotted lines respectively. Scale bar: 5µm.* ***(C and D)*** *Quantification of cells number for localization of* ***(C)*** *SMN (in Cajal Bodies [CBs] or Gems, at the periphery of the nucleolus or mixed localization) and* ***(D)*** *RNAP1 (inside the nucleolus, outside the nucleolus or mixed localization) at different times PUVI in cells depleted of the endogenous SMN, by induction of the sh6-SMN RNA, and expressing GFP-SMN, GFP-SMN*$^{E134K}$ *or GFP-SMN*$^{Y272C}$ *.*

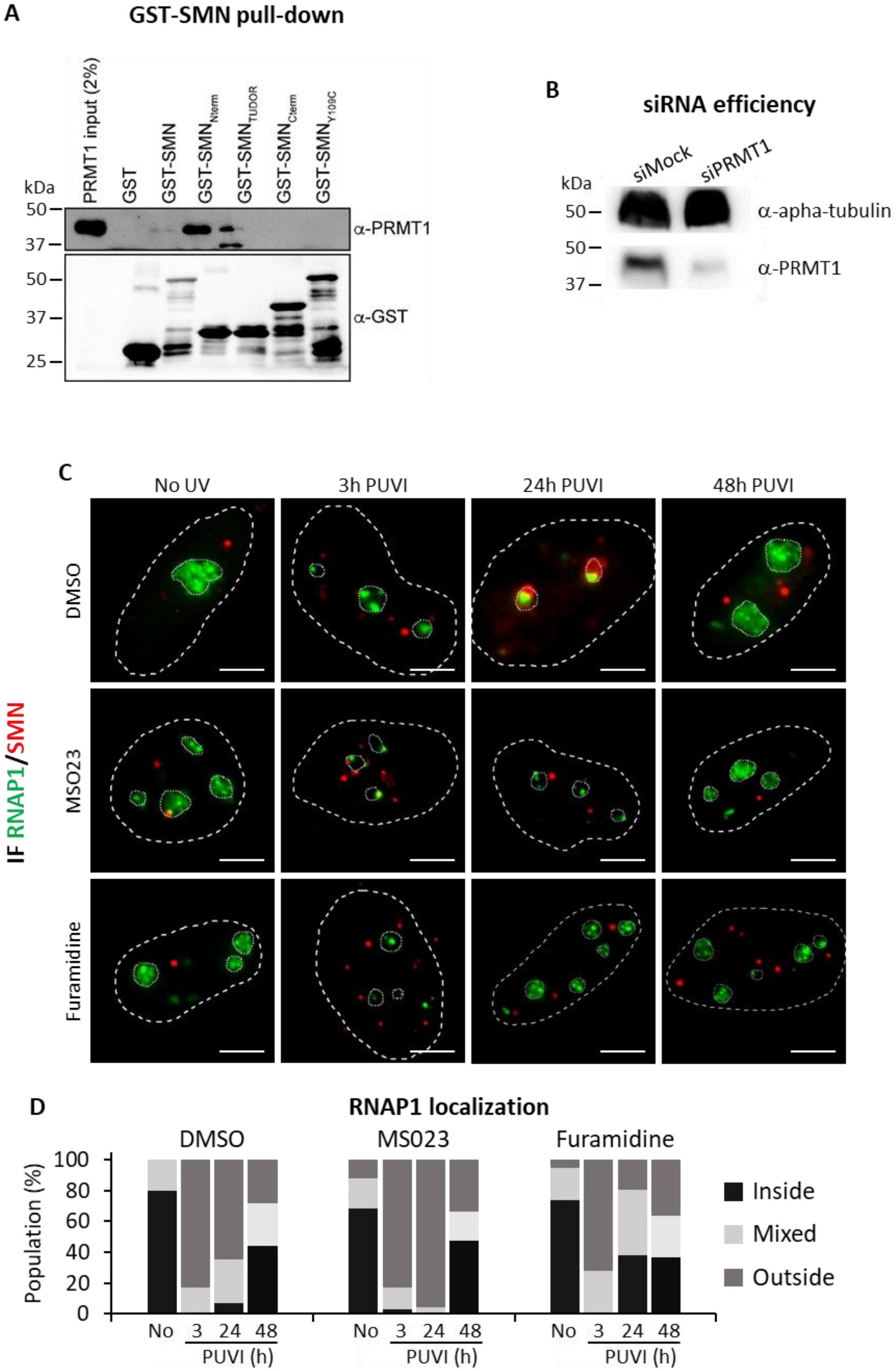
A
GST-SMN pull-down
PRMT1 input (2%)
GST
GST-SMN
GST-SMN Nterm
GST-SMN TUDOR
GST-SMN Cterm
GST-SMN Y109C
kDa
50
37
50
37
25
α-PRMT1
α-GST
B
siRNA efficiency
siMock
siPRMT1
kDa
50
50
37
α-apha-tubulin
α-PRMT1
C
No UV
3h PUVI
24h PUVI
48h PUVI
DMSO
MSO23
Furamidine
IF RNAP1/SMN
D
RNAP1 localization
DMSO
MS023
Furamidine
Population (%)
100
80
60
40
20
0
No
3
24
48
PUVI (h)
Inside
Mixed
Outside

***Figure S12: Relation between PRMT1 and SMN.***

***(A)*** *GST pull-down assay using purified recombinant SMN protein with different truncations of the protein and cellular extracts.* ***(B)*** *Western Blot on whole cell extracts of MRC5 cells treated with siPRMT1.* ***(C)*** *Representative microscopy images of IF assay showing the localization of SMN (red) and RNAP1 (green) in MRC5 cells treated with DMSO, MS023 or Furamidine followed by 16J/m² UV-C irradiation. Nuclei and nucleoli are indicated by dashed lines and dotted lines respectively. Scale bar: 5µm.* ***(D)*** *Quantification of cells number for localization of RNAP1 (inside the nucleolus, outside the nucleolus or mixed localization) at different times PUVI in cells treated with DMSO, MS023 or Furamidine.*

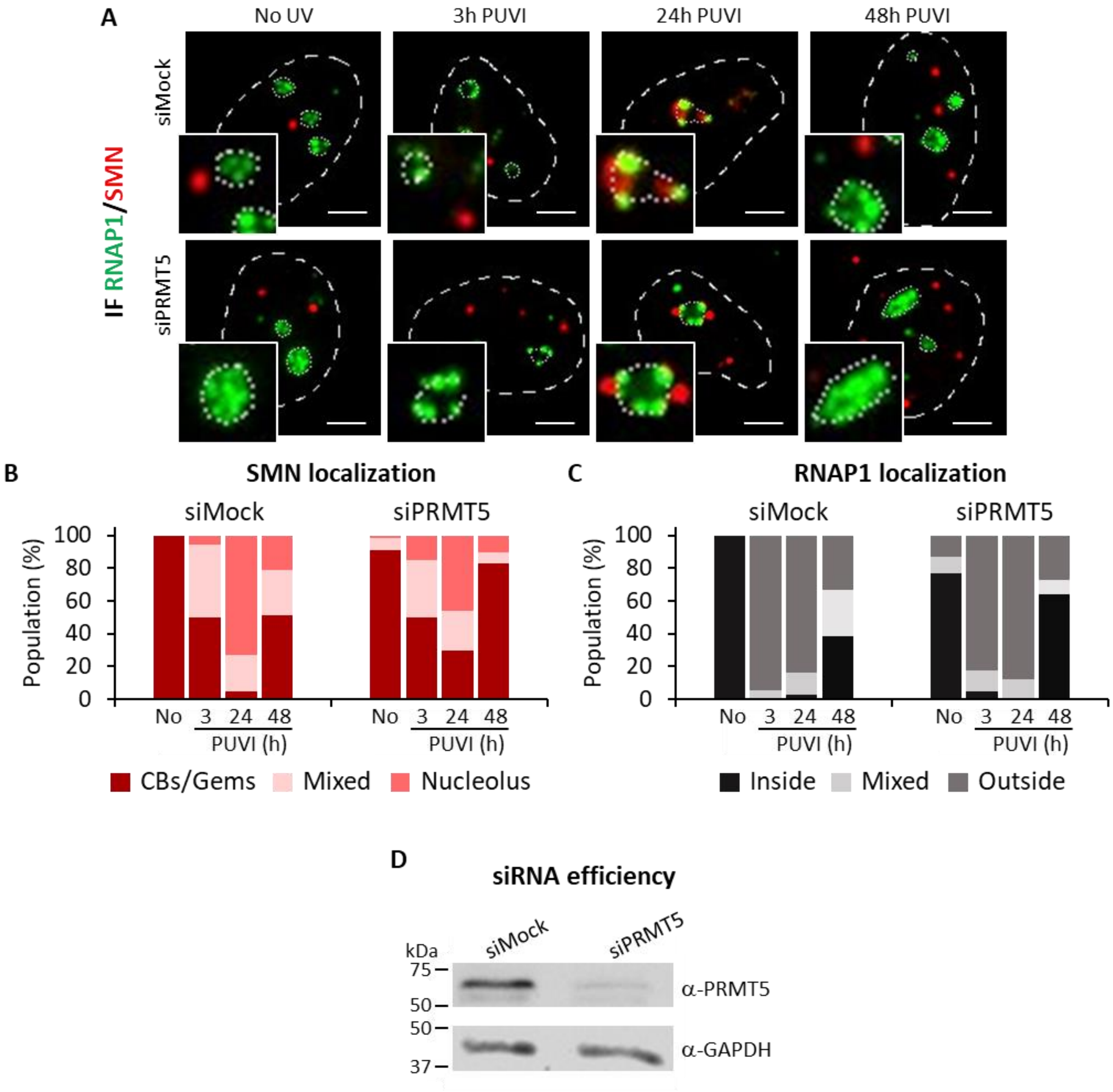


**Figure S13: SMN shuttling is PRMT5-independent**

***(A)*** *Representative microscopy images of immunofluorescence (IF) assay showing the localization of SMN (red) and FBL (green) at different times PUVI in cells transfected with siMock or siPRMT5 pool. Nuclei and nucleoli are indicated by dashed lines and dotted lines respectively. Scale bar: 5µm.* ***(B and C)*** *Quantification of cells number for localization of* ***(B)*** *SMN (in Cajal Bodies [CBs] or Gems, at the periphery of the nucleolus or mixed localization) and* ***(C)*** *FBL (inside the nucleolus, outside the nucleolus or mixed localization) at different times PUVI in cells transfected with siMock or siPRMT5 pool.* ***(B)*** *Western Blot on whole cell extracts of MRC5 cells treated with siPRMT5.*

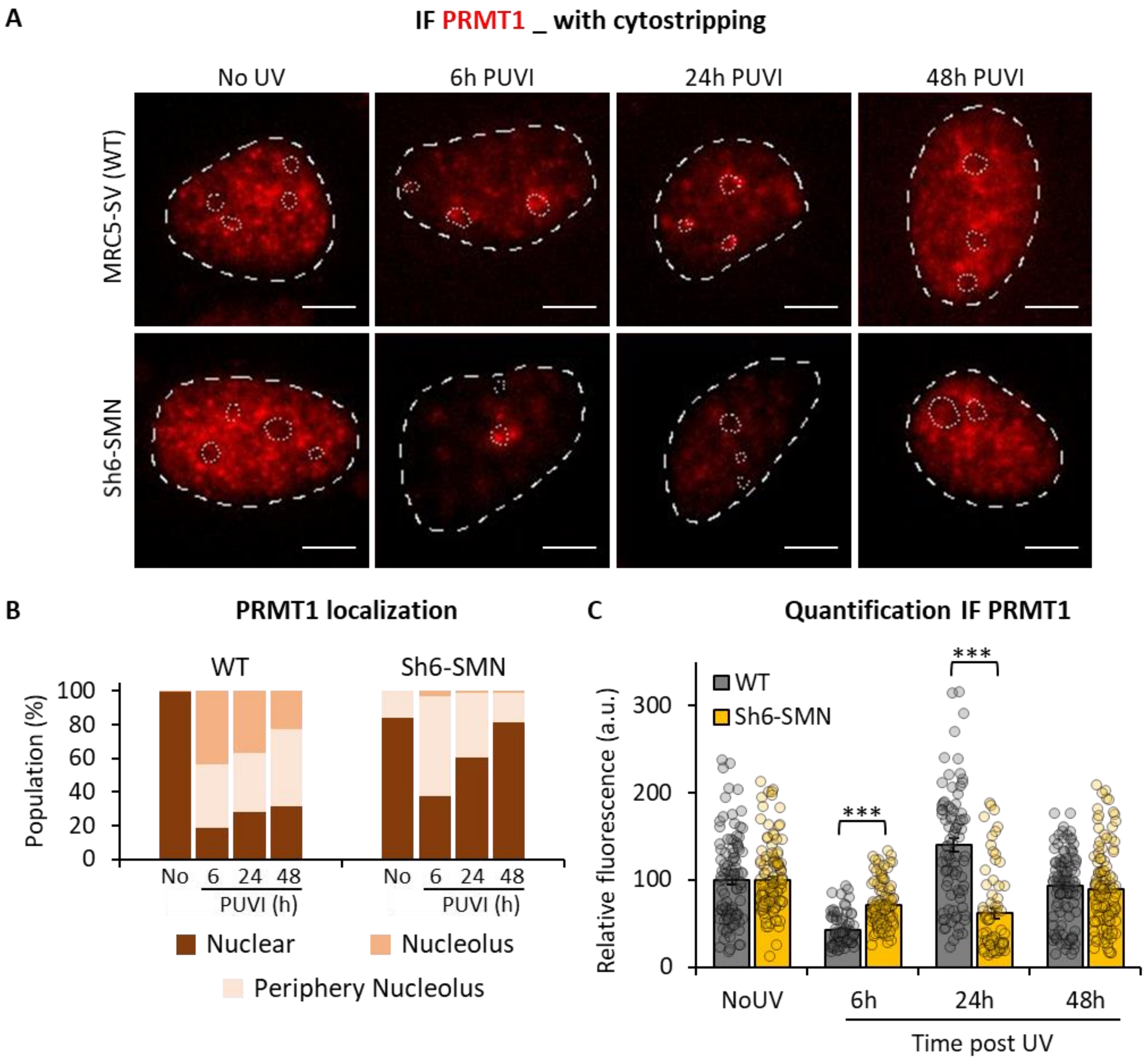


**Figure S14: PRMT1 localization and expression after UV irradiation in the presence or absence of SMN**

***(A)*** *Representative microscopy images of IF with cytostripping showing the localization of PRMT1 (red) in MRC5 and Sh6-SMN cells after UV-C irradiation. Nuclei and nucleoli are indicated by dashed lines and dotted lines respectively. Scale bar: 5µm.* ***(B)*** *Quantification of cells number for localization of PRMT1 (pan nuclear, at the periphery of the nucleolus or inside the nucleolus) at different times PUVI in WT or SMN depleted cells (Sh6-SMN).* ***(C)*** *Quantification of fluorescent signal in the nucleus from the IF with cytostripping against PRMT1. Error bars represent the standard error of the mean (SEM) obtained from at least 60 cells. P-value of Student's test compared to No UV condition. ***<0.001.*

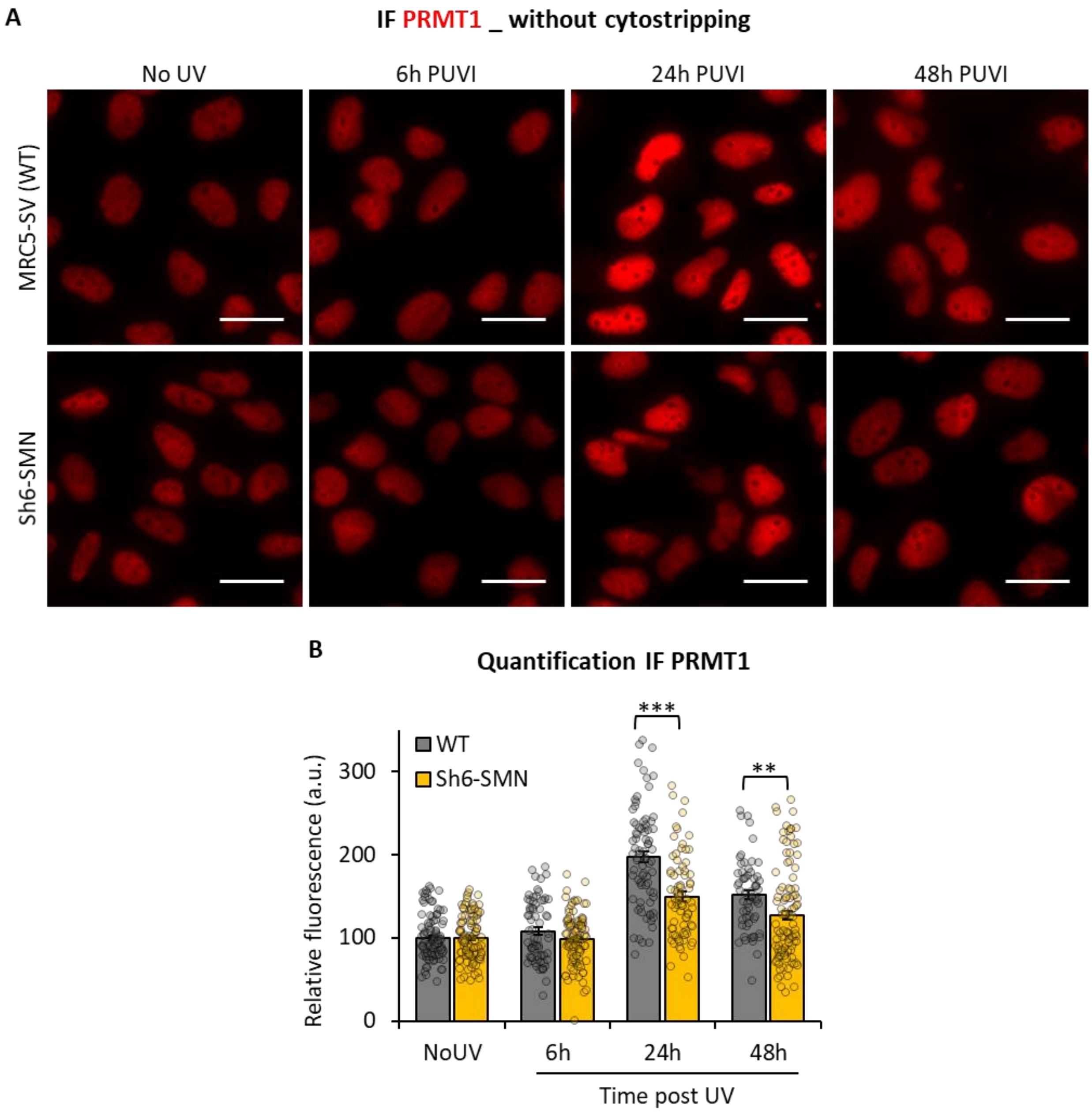


**Figure S14: PRMT1 localization and expression after UV irradiation in the presence or absence of SMN (without cytostripping)**

***(A)*** *Representative microscopy images of IF without cytostripping showing the localization of PRMT1 (red) in MRC5 and Sh6-SMN cells after UV-C irradiation. Nuclei and nucleoli are indicated by dashed lines and dotted lines respectively. Scale bar: 5µm* ***(C)*** *Quantification of fluorescent signal in the nucleus from the IF without cytostripping against PRMT1. Error bars represent the standard error of the mean (SEM) obtained from at least 60 cells. P-value of Student's test compared to No UV condition. **<0.05, ***<0.001.*

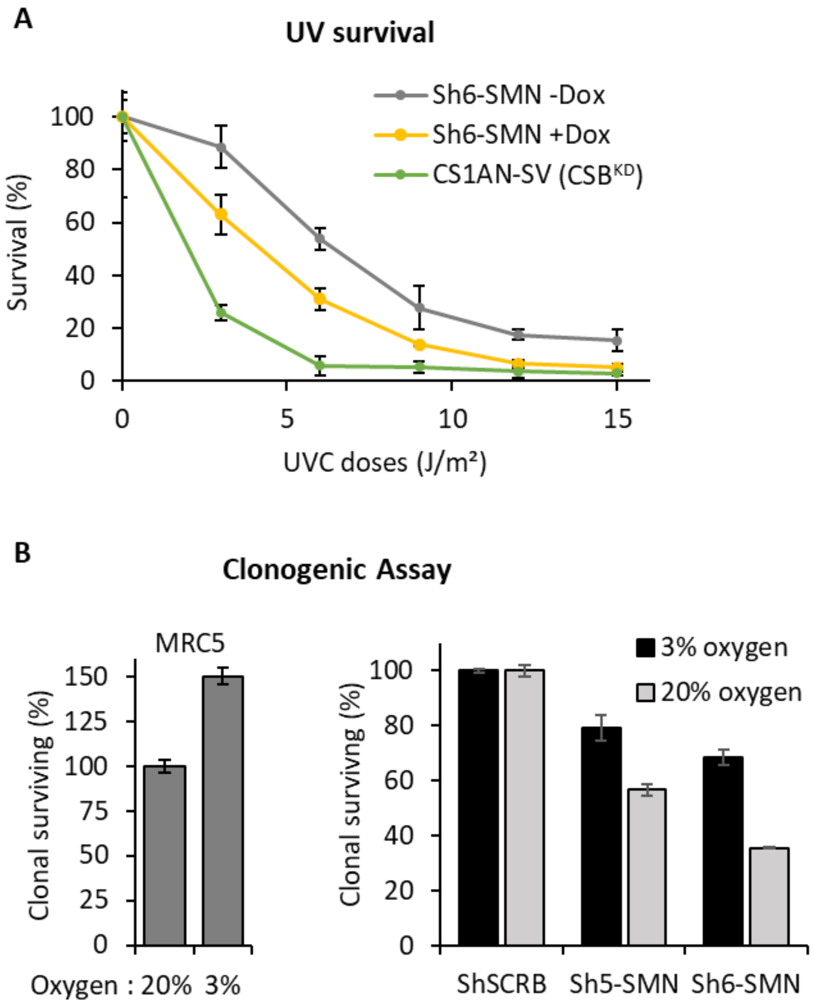


**Figure 8: SMN-deficient cells are sensitive to DNA damage**

***(A)*** *Survival curve as determined by the colony-forming ability to UV-C of cells expressing (+Dox) or not (-Dox) Sh6-SMN.* ***(B)*** *Clonogenic assay of cells cultured with different quantities of oxygen (3% or 20%) in the presence (MRC5 or ShSCRB) or absence of SMN (Sh5-SMN and Sh6-SMN).*

### MATERIALS AND METHODS supplemental

#### Cells culture

Primary fibroblast cells from unaffected (C5RO) and CSB-deficient patients (CS1AN) were cultured in DMEM supplemented with 10% FBS and 1% P/S. SMA type I patients (GM00232) fibroblast cell lines were obtained from Coriell Cell Repositories and cultured in MEM supplemented with 15% non-inactivated FBS, 1% non-essential amino acids and 1% P/S. All primary fibroblasts are incubated at 37°C with 3% O2 and 5% CO2.

Human iPSC-derived motoneurons (hMNs) were generated as previously described by [1] Human iPSCs were dissociated with TrypLE (Gibco) and resuspended in motoneuron medium (MNB) containing DMEM-F12 Glutamax/Neurobasal (1:1 ratio; Gibco), N2 supplement (Gibco), B27 without vitamin A supplement (Gibco), β-mercaptoethanol (0,1%; Gibco), Penicillin/Streptomycin (0,1%; Gibco), supplemented with small molecules including ascorbic acid (0.5 µM; Sigma-Aldrich), SB431542 (20 µM; TOCRIS-BioTechne, Minneapolis, MN, USA), LDN193189 (0.2 µM; Miltenyi Biotec), CHIR99021 (3 µM; Miltenyi Biotec) and Y-27632 (10 µM; STEMCELLS Technologies, Vancouver, Canada) for the first day of differentiation. Cells were seeded in suspension into 6 well plate (Dutscher, Bernolsheim, France) treated with anti-adherence rinsing solution (Stemcell, Technologies) to form spheroids. During the entire culture process, small molecules were added at different time points including retinoic acid (0.1 µM RA; Sigma-Aldrich), Smoothened Agonist (0.5 µM SAG; STEMCELLS Technologies), Brain-Derived Neurotrophic Factor (10 ng/mL BDNF; PreproTech, Rocky Hill, NJ, USA) and γ-secretase inhibitor (10 µM DAPT; STEMCELLS Technologies). Then, spheroids were dissociated at DIV 10 (days in vitro) with trypLE to obtained hMNs progenitors that were plate on glass coverslips coated with poly-L-ornithine (Sigma-Aldrich) and laminin at 1-2µg/ml (Sigma-Aldrich) or dispensed into cryovials and freezed using CryoMed Controlled-Rate Freezer (Thermo Fisher Scientific). hMNs were maintain in culture for maturation in MNB supplemented with BDNF and GDNF.

#### Construction and expression of GFP-SMN and Cherry-SMN fusion protein

The GFP-SMN WT and GFP-SMN E134K were previously described in [2]. The GFP-SMN Y272C was generated using standard PCR-based site-directed mutagenesis. For mCherry-SMN WT, SMN cDNA was inserted in mCherry-N1.

Cherry-SMN WT and GFP-SMN WT, E134K and Y272C stably expressing cell lines were produced by transfecting the plasmid in MRC5-SVcells containing sh6-SMN. FuGENE 6 Transfection Reagent (Promega) was used according to the manufacturer' protocol. The selection was performed with G418 at 200 µg/mL.

**Treatment**

RNAP1 transcription inhibition has been achieved by incubation in medium containing Cordycepin at 50µg/mL. Resumption of transcription has been obtained by replacement of Cordycepin medium with normal medium after two washes with PBS. No treated cells were used as control.

The stock of KBr03 is freshly diluted at 80mM in warm DMEM. After one wash with PBS, cells are incubated one hour with KBr03. After this incubation, cells were washed two time with PBS and incubated with medium at 37°C with 5% C02 for different period of time.

**Transfection of small interfering RNAs (siRNAs)**

The small interfering RNA (siRNAs) used in this study are: siPRMT5, GGCCAUCUAUAAAUGUCUG (10mM), SiCOILIN #02, GAGAGAACCUGGGAAAUU (10nM), SiCOILIN 17, CGGAGUGUGCUGCGGGUUU (10nM), siFBL 05 GGUCGAGGCGGAGGCUUUA (10nM), siFBL 06, AAUGGUGGAUGUGAUCUUU (10nM).The final concentration used for each siRNA is indicated in parentheses.

**GST-SMN purification and GST pull-downs**

SMN (full-length or truncated) cDNA was cloned in pGEX6P1 between *BamHI* and *XhoI* sites and transformed in BL21 (DE3) cells (200131; Agilent). Single colonies were grown overnight in 2.5 mL LB broth, scaled up to 250 mL, grown at 37 °C until density at $OD_{600}$ reached 0.6, then GST or GST-SMN were induced with 0.2 mM IPTG overnight. The next day, cells were collected by centrifugation and resuspended in 10 mL lysis buffer (50mM Tris pH 8.0, 150mM NaCl, 0.05% NP40, supplemented with PIC). While working on ice, cells were briefly sonicated and extracts clarified by centrifugation. Recombinant proteins were then purified using Glutathione-sepharose beads (tumbling at 4 °C overnight), washed extensively with lysis buffer, and released from the beads using elution buffer (100mM Tris pH 8.0, 10% Glycerol, 15mg/mL reduced glutathione).

GST pull-downs were performed in 600 µL TAP buffer (50mM Tris pH 7.5, 200mM NaCl, 0.1% Triton-X100, and 10% glycerol supplemented with PIC) with 5µg GST and 85µL HEK293T whole cell lysate (1 x 100 mm plate lysed in 1 mL TAP buffer). A 10% input (8.5µL in 20µL Laemmli sample buffer) was set aside. Samples were incubated 2-3h at 4°C with rotation, then 25µL Glutathione-sepharose beads were added for 1h. Finally, the beads were washed 4 times with 1mL TAP buffer and finally resuspended in 20µL Laemmli sample buffer before immunoblotting analyses.

For time course experiments following UV-induced DNA damage, MRC5-SV cells were lysed in Pierce IP lysis buffer (Thermo 87787) and 17µg of proteins (amount available per pull-down) were used as above.

**Recovery of RNA synthesis (RRS) assay**

Cells were grown on coverslips. RNA detection was done using a Click-iT RNA Alexa Fluor Imaging kit (Invitrogen, C10330), according to the manufacturer's instructions. Briefly, cells were UV-C irradiated (10 J/m²) and incubated for 3 or 24 h at 37°C. Then, cells were incubated for 2 hours with 5-ethynyl uridine (EU). After fixation and permeabilization, cells were incubated for 30min with the Click-iT reaction cocktail containing Alexa Fluor Azide 594. After washing, the coverslips were mounted with Vectashield (Vector). The average fluorescence intensity per nucleus was estimated after background subtraction using ImageJ and normalized to not treated cells. At least 50 cells were images for each condition of each cell lines.

**Unscheduled DNA synthesis (UDS or TCR-UDS)**.

Cells were grown on coverslips. After local irradiation, cells were incubated for 3 or 8 hours (UDS and TCR-UDS respectively) with 20µM of 5-ethynyl-2'-deoxyuridine (EdU), fixed with 4% PFA for 15min at 37°C and permeabilized with PBS and 0.5% Triton X-100 for 20min. Then, cells were blocked with PBS+ for 30min and subsequently incubated for 1h at RT with mouse monoclonal anti-yH2AX antibody (Ser139 [Upstate, clone JBW301]) 1:500 diluted in PBS+. After extensive washes with PBS containing 0.5% Triton X100, cells were incubated for 45min at RT with secondary antibodies conjugated with Alexa Fluor 594. Next, cells were washed several times and then incubated for 30 min with the Click-iT reaction cocktail containing Alexa Fluor Azide 488 (Invitrogen, C10337). After washing, the coverslips were mounted with Vectashield containing DAPI (Vector). Images of the cells were obtained with the same microscopy system and constant acquisition parameters.

Images were analyzed as follows using ImageJ and a circle of constant size for all images: (i) the background signal was estimated in the nucleus (avoiding the damage, nucleoli and other non-specific signal) and subtracted, (ii) the locally damaged area was defined by using the yH2AX staining, (iii) the average fluorescence correlated to the EdU incorporation was then measured and thus an estimate of DNA synthesis after repair was obtained. For UDS experiment, at least 30 cells were imaged for each condition. For TCR-UDS, at least 20 cells were imaged for each condition.

**Primary Antibodies**

The following primary antibodies were used:

| **Antibody against** | **Manufacturer** | **Catalog Nr** | **Source** | **WB** |
|---|---|---|---|---|
| Cherry | Abcam | Ab167453 | Rabbit | 1/2000 |
| GFP | Sigma-aldrich | SAB4701015 | Rabbit | 1/1000 |

| | | | | |
|---|---|---|---|---|
| GST | Abcam | ab3416 | Rabbit | 1/3000 |
| PRMT5 | Upstate | 07-405 | Rabbit | 1/1000 |
